# Long-term evolution reveals the role of the circadian cycle in the environmental adaptation of cyanobacteria

**DOI:** 10.1101/2024.03.12.584591

**Authors:** Alfonso Mendaña, María Santos-Merino, Raquel Gutiérrez-Lanza, Marina Domínguez-Quintero, Juan Manuel Medina, Ana González-Guerra, Víctor Campa, Magaly Ducos-Galand, Rocío López-Igual, Daniel C. Volke, Muriel Gugger, Pablo I. Nikel, Didier Mazel, Fernando de la Cruz, Raúl Fernández-López

## Abstract

Circadian clocks synchronize internal cellular states with diurnal rhythms. Widespread in bacteria and eukaryotes, they regulate a variety of physiological processes, from hormone secretion in animals to carbon fixation in photosynthetic organisms. The adaptive role of circadian clocks is assumed to stem from their ability to anticipate environmental change, yet their impact on ecological adaptation remains unclear. Here, we use experimental evolution to study the interplay between fitness and circadian regulation in the model cyanobacterium *Synechococcus elongatus* PCC 7942. After 1,200 generations under continuous, high-intensity illumination, we obtained a strain that grew six times faster than its ancestral counterpart. Genome sequencing revealed three mutations fixed in the population, two of which replicated the fast-growing phenotype in the wild-type. A deletion in SasA, a key circadian regulator, was essential for fast growth. Transcriptomic and metabolomic analyses revealed that this mutation perturbed the rhythmicity of the cycle, while simultaneously locking the cell in a transcriptomic response to high intensity illumination. A comparison with another fast-growing isolate, UTEX 2973, showed convergent transcriptomic states despite different driving mutations. Our results indicate that the clock acts not only as a timekeeping device, but also as an adaptive mechanism to optimize growth across diverse ecological conditions.

---

Initially considered to be too simple to sustain a circadian clock, cyanobacteria have emerged as one of the best-studied models of circadian regulation. Key players in the carbon and oxygen cycles on earth, cyanobacteria employ photosynthesis to produce ATP and reducing power. To avoid the generation of lethal reactive oxygen species, cyanobacteria must carefully balance their metabolism with external light, thus their physiology is heavily influenced by circadian rhythms^1^. The cyanobacterial clock is based on a post-transcriptional oscillator (PTO) formed by the products of the *kaiABC* cluster^2,3,4,5,1^. Fluctuations in the PTO are converted into transcriptional cycles through RpaA, the master transcriptional regulator of the circadian cycle^6,7^. Genes controlled by RpaA can be divided into two classes^8^. Class I genes are activated by RpaA and achieve a maximum at dusk, while Class II are repressed by RpaA, and peak at dawn^6,9,7^. In the model cyanobacterium *Synechococcus elongatus* PCC 7942, 64% of the genes are under circadian transcriptional regulation^8,10^.

Maintaining energy levels through the regulation of glycogen metabolism has been the classical function assigned to the clock^11,12,13^. Yet in PCC 7942 a wide range of physiological functions, such as stress responses, natural competence and cell division, are regulated in a circadian fashion, suggesting that the clock may play roles beyond energetic balance^14,15,16^. Despite our knowledge on its molecular mechanisms, the ecological role of circadian regulation remains elusive. While the core components of the clock are highly conserved across cyanobacteria, its physiological relevance varies between species^17,18^. Marine picocyanobacteria, such as *Prochlorococcus*, contain a simpler mechanism lacking key features of a *bona fide* clock^19^. Other model species, like *Synechocystis*, exhibit less marked circadian oscillations^20^; and in *Cyanothece* sp. ATCC 51142, ultradian rhythms longer than 24 hours have been observed^21^. Overall, the reasons for these differences are unclear. It has been shown that similar cyanobacteria exhibit substantial divergence in growth rates and environmental preferences, even among isolates of the same species^22^. In *S.elongatus*, for example, model strains like PCC 7942 and PCC 6301 tolerate moderate light intensities and show optimal doubling times of approximately 6-8 h. Yet, UTEX 2973, a strain that is 99.99% identical at the DNA level, grows optimally at much higher light intensities, with doubling times as short as 2 h^23^.

To identify the genetic basis of this phenotypic plasticity, we subjected PCC 7942 to a long-term evolution experiment (LTE). The strain was grown under continuous, high intensity illumination for 1,200 generations. At the end of the LTE, the evolved population grew six times faster than its ancestral counterpart. Genetic, transcriptomic and metabolomic analyses revealed that adaptation to fast growth involved profound alterations in the circadian cycle. A comparison between the transcriptomes of our evolved strain and UTEX 2973 showed that both strains adapted to growth under high light intensities through convergent transcriptional strategies, yet the driving mutations were different. Altogether, our results demonstrate that mutations in circadian control have drastic effects on the cyanobacterial phenotype and are key for *S. elongatus* to adapt to different environmental light regimes.

## RESULTS

### LONG-TERM EVOLUTION

PCC 7942, a strain with a planktonic phenotype, unable to form biofilms, was chosen for a long-term evolution experiment under serial passage^24^. A master culture of PCC 7942 was retrieved from the Pasteur Culture Collection to serve as the initial population. During the evolution experiment, cells were grown in BG11 medium at 41°C, 1,300 µmol photons m^-2^ s^-1^, under continuous illumination and constant bubbling of 5% CO_2_. Culture dilutions were performed every 24 generations, starting with a 1/130 dilution that ensured an effective population size of at least 10^9^ cells. After 1,248 generations, the growth rate of the evolved population, named C11, was compared to the ancestral strain. Results showed that, under conditions of high light intensity (HL, 983 µmol photons m^-2^ s^-1^), high temperature (HT, 41°C), and high CO_2_ levels (HC, 3%), the generation time of C11 was 1.6 h, representing a 654% increase compared to the wild-type (wt) (Figure 1A, left). This fitness gain, however, was contingent on the environmental conditions. Reducing the illumination to 120 µmol photons m^-2^ s^-1^ (LL) or lowering the temperature to 30°C (LT) resulted in a decrease in the fitness of C11 (Figure 1B). CO_2_ levels showed the largest impact on the doubling time. When C11 was grown under atmospheric CO_2_ concentrations (LC, 0.04%), its growth rate decreased 6- fold, regardless of light intensity (Figure 1A, right). Thus, at low CO_2_ levels the wt grew faster than C11 in all conditions (Supplementary Figure 1). Results thus showed that PCC 7942 adapted quickly to the conditions of the LTEE, increasing its fitness by more than 600%, yet its adaptation was contingent on the environmental levels of CO_2,_ light, and temperature.

**Figure 1.-.**
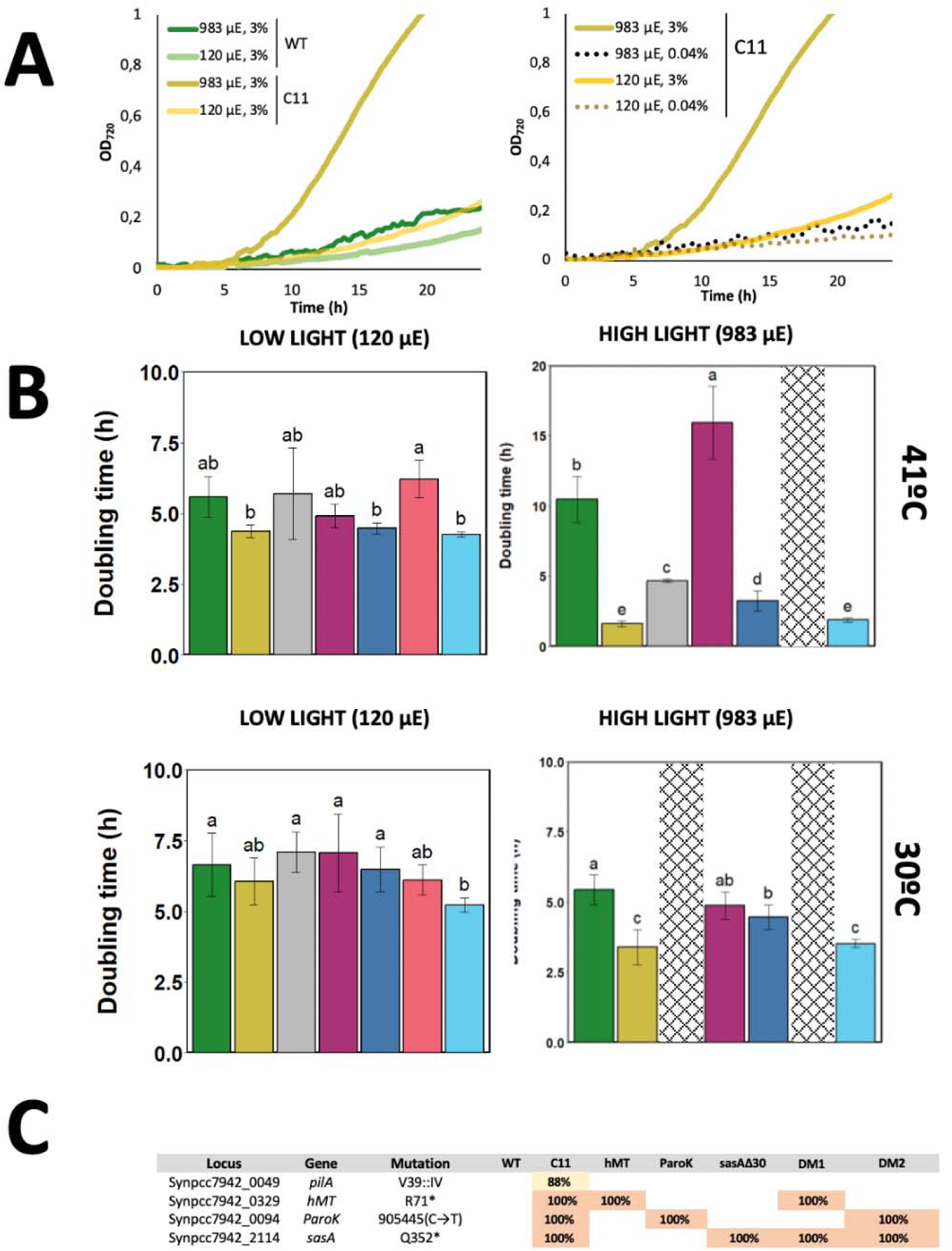
Growth rates of S. elongatus PCC 7942 ancestral and evolved strains in different conditions. (A) Growth curves of the ancestral (wt) and evolved (C11) strains monitored as the OD_750_ (y axis) along time (x axis). The panel on the left shows the growth advantage of C11 at high light intensity (HL, 983 µmol photons m^-2^ s^-1^, dark yellow line) compared to the growth curves of C11 at moderately low light intensity (LL, 120 µmol photons m^-2^ s^-1^ light yellow line) and the wt at HL (dark green line) and LL (light green line) intensities. All growth curves on this panel were carried out at 41°C (HT) and 3% CO_2_ (HC) saturation. The right panel shows the growth advantage at HL intensity (dark yellow line) disappearing when the culture is grown at atmospheric (0.04%, LC) CO_2_ concentrations (dashed lines). (B) Growth rates, expressed as the doubling time, of C11, wt and point mutants at different temperatures (41°C upper, 30°C lower) and light intensities (120 µmol photons m^-2^ s^-1^, LL, left; 983 µmol photons m^-2^ s^-1^, HL, right). Bars indicate the mean and standard deviation of at least four independent replicates. Dashed bars represent conditions for which no growth was observed. Bars with the same letter within the same condition do not differ statistically (p>0.05) by Tukey’s multiple comparison test. (C) Location of the mutations fixed or nearly fixed (>88% reads) in C11. Percentages indicate the fixation index of each mutation in the evolved strain (C11) and point mutants, named after their respective mutations. DM1 and DM2 indicate, respectively, the *hMT*/*sasA*Δ30 and *sasA*Δ30/P*aroK* double mutants.

### RECONSTRUCTION OF THE PHENOTYPE IN THE WILD TYPE

To identify the genetic basis of adaptation, we sequenced the genome of C11 and compared it with its ancestral strain. A frozen stock of the evolving population obtained at generation 824 (G824) was also introduced in the analysis (Supplementary Table 1). When the genomes of C11 and PCC 7942 were compared, we identified four mutations that were fixed or nearly fixed (present in >80% of the reads) in C11 (Figure 1C). A mutation in *pilA1* (Synpcc7942_0048), a gene required for natural competence in cyanobacteria, was found in 88% of the C11 population. This mutation caused the loss of natural competence (Supplementary Figure 2), a phenotype often observed in other model cyanobacterial strains^22,25^. Three additional mutations were fixed in the C11 population. The first one caused an early stop codon in the gene Synpcc7942_0329, coding for a putative class I SAM-dependent methyl transferase (Refseq: WP_011377536). We named this mutation *hMT,* from *h*ypothetical *M*ethyl *T*ransferase. Homologs in *Synechocystis* sp. PCC 6803 and *Synechococcus* sp. PCC 7002 (CpcM) have been involved in the posttranslational methylation of phycobiliproteins, contributing to the efficiency of energy transfer in the photosynthetic chain^26^. This mutation produces a truncation of the polypeptide at R71 which, given the length of the wt protein, is likely to result in loss of function. The second mutation (P*aroK*) involves a C→T transition in the promoter region of *aroK* (Synpcc7942_0894), a gene encoding the shikimate kinase. AroK catalyzes the fifth reaction in the chorismate synthesis pathway^27^. Finally, the third mutation (*sasA*Δ30) causes a premature stop codon in *sasA* (Synpcc7942_2114), a gene involved in circadian regulation. SasA is a histidine-kinase that phosphorylates RpaA, the master regulator of circadian promoters^28,6^. The *sasA*Δ30 mutation in C11 truncates SasA with a premature stop codon at Q352, thus generating a 30 amino acid deletion at the C-terminus of the protein, which contains the histidine kinase domain. Among these mutations, both P*aroK* and *sasA*Δ30 were already present in the population at G842 (Supplementary Table 1), whereas *hMT* emerged later in the evolutionary process.

To test the impact of these mutations, we introduced them in the genome of PCC 7942, replacing wt alleles with their mutant counterparts (see Materials and Methods). The growth rate of each individual mutant was measured under different environmental conditions and compared to the wt (Figure 1B). Although both *sasA*Δ30 and *hMT* grew significantly faster than the wt in the conditions of the LTE, their growth rate was smaller than C11. The introduction of the ParoK mutation alone did not improve growth in any of the conditions tested. We thus introduced combinations of these mutations in PCC 7942. A combination of *hMT* with *sasA*Δ30 was found to be lethal under HL conditions, and all our attempts to generate a triple mutant were unsuccessful. However, as shown in Figure 1B, the combination of P*aroK* and *sasA*Δ30 resulted in growth rates like C11. As in the evolved population, this growth advantage was only observed in HL, HT and HC conditions.

### THE EVOLVED STRAIN IS LOCKED IN A TRANSCRIPTIONAL RESPONSE TO HIGH LIGHT

To unveil the causes of fast growth, we conducted transcriptomic analyses of C11 and the wt under HL and LL conditions. Results are detailed in Supplementary Table 2. We first identified genes responding to light intensity by calculating the log2 fold-change between LL and HL in the wt (Figure 2A). Alongside these results, we plotted the log2 fold-change between the wt and C11, both at LL and HL. As shown in Figure 2A, many genes repressed by HL in the wt were constitutively downregulated in C11 (arrows in Figure 2A). Among them, we identified hydrogenases (Figure 2B), genes involved in glycogen metabolism (Figure 2C), and regulatory genes (Figure 2D). A subset of genes that, in the wt, increased their transcription in response to HL were also constitutively induced in C11 (Supplementary Figure 3). Thus, the constitutive transcriptome of C11 mirrored the high-light response exhibited by the wt.

**Figure 2.-.**
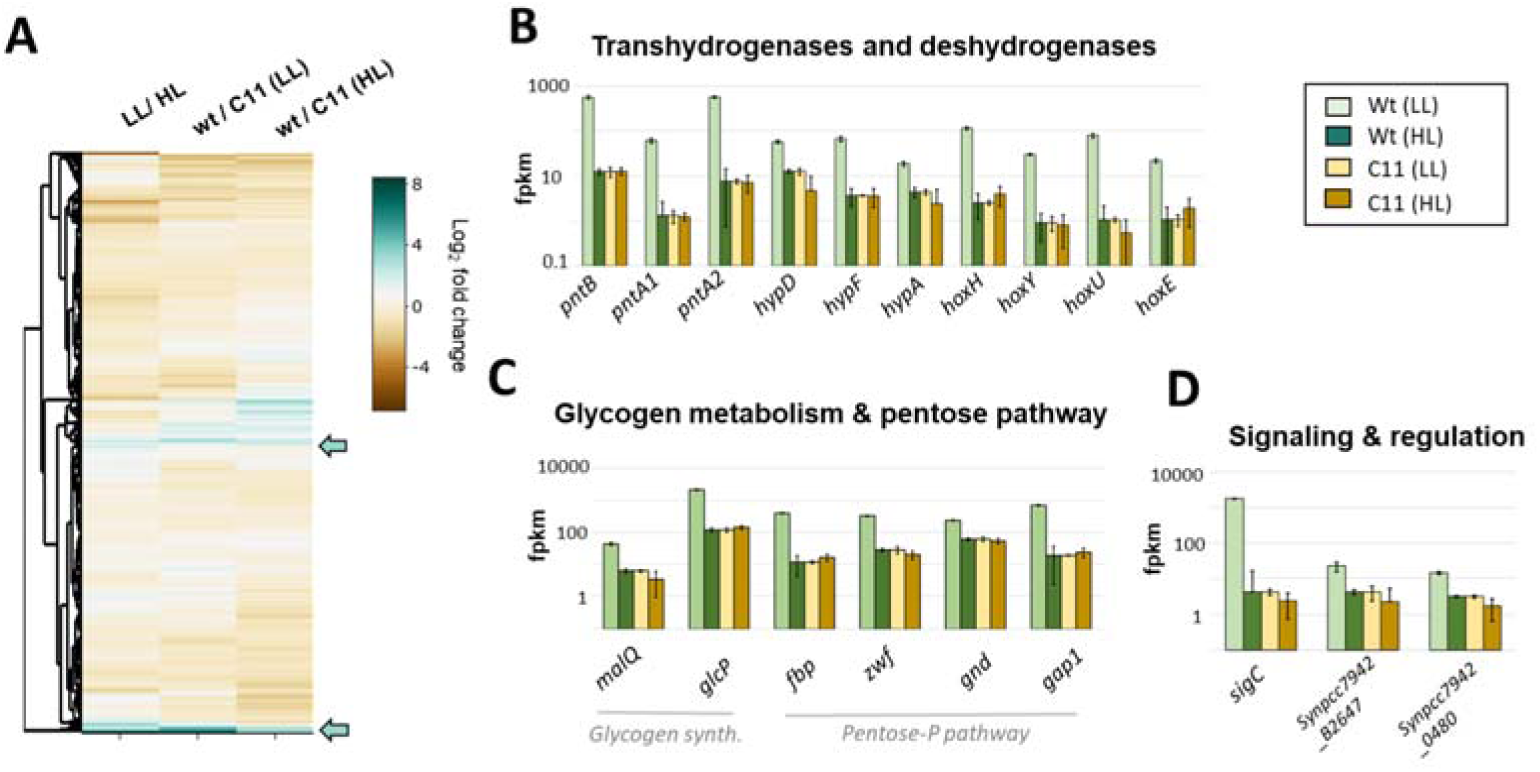
C11 transcriptome is locked in a high-light response mode. (A) Heatmap showing the changes in gene expression, indicated as Log_2_ of the fold change in FPKM, in the wt and C11 at different light intensities. The leftmost lane corresponds to fold changes experienced by the wt when grown in moderately low light (LL, 120 µmol photons m^-2^ s^-1^) compared to high light (HL, 983 µmol photons m^-2^ s^-1^). The central lane corresponds to the changes observed in C11 compared to wt when both strains were grown under LL intensities. The rightmost lane shows the same comparison, but measured when C11 and the wt were grown under HL intensity. Arrows indicate genes showing decreased transcription in HL in the wt that are constitutively repressed in C11. Charts B, C, and D on the right, show transcriptional levels, expressed as the average FPKM and the standard deviation of three independent replicates, of genes repressed in HL in the wild type, which are constitutively shut down in C11. Genes shown are involved in hydrogenases (B), glycogen synthesis and the pentose phosphate pathway (C) and signaling and regulation (B).

Since transcriptomic changes involved key metabolic pathways, we carried out a metabolomic comparison between C11 and the wt. As described in Materials and Methods, 112 metabolites were measured following the protocol of Prasannan et al.^29^ Results, shown in Supplementary Table 3, demonstrated that C11 exhibited increased levels of many intermediates of the Calvin-Benson-Bassham (CBB) and the Tricarboxylic Acid (TCA) cycles (Figure 3). The most significant increases were observed in fructose 6-phosphate and fructose 1,6-bisphosphate. These changes were concomitant to alterations in the expression of key enzymes of the CBB and TCA pathways, as judged from transcriptomic data (Supplementary Table 2). PCC 7942 contains two isoforms of the fructose 1,6-bisphosphatase: *fbp* and *glpX*^30^. In C11, the expression of the former was downregulated, while the latter was moderately increased. Similarly, C11 exhibited a downregulation of the *gap1* gene, which encodes an NAD-dependent glyceraldehyde-3P-dehydrogenase. However, the expression of *gap2* gene, which encodes an NADP-dependent isoform, was unchanged. In *Synechocystis*, *gap1* is involved in catabolism, while *gap2* has anabolic functions^31^. Altogether, metabolomic and transcriptomic data showed a significant alteration of carbon metabolism, with a global repression of glycogen catabolism and the pentose phosphate pathways.

**Figure 3.-.**
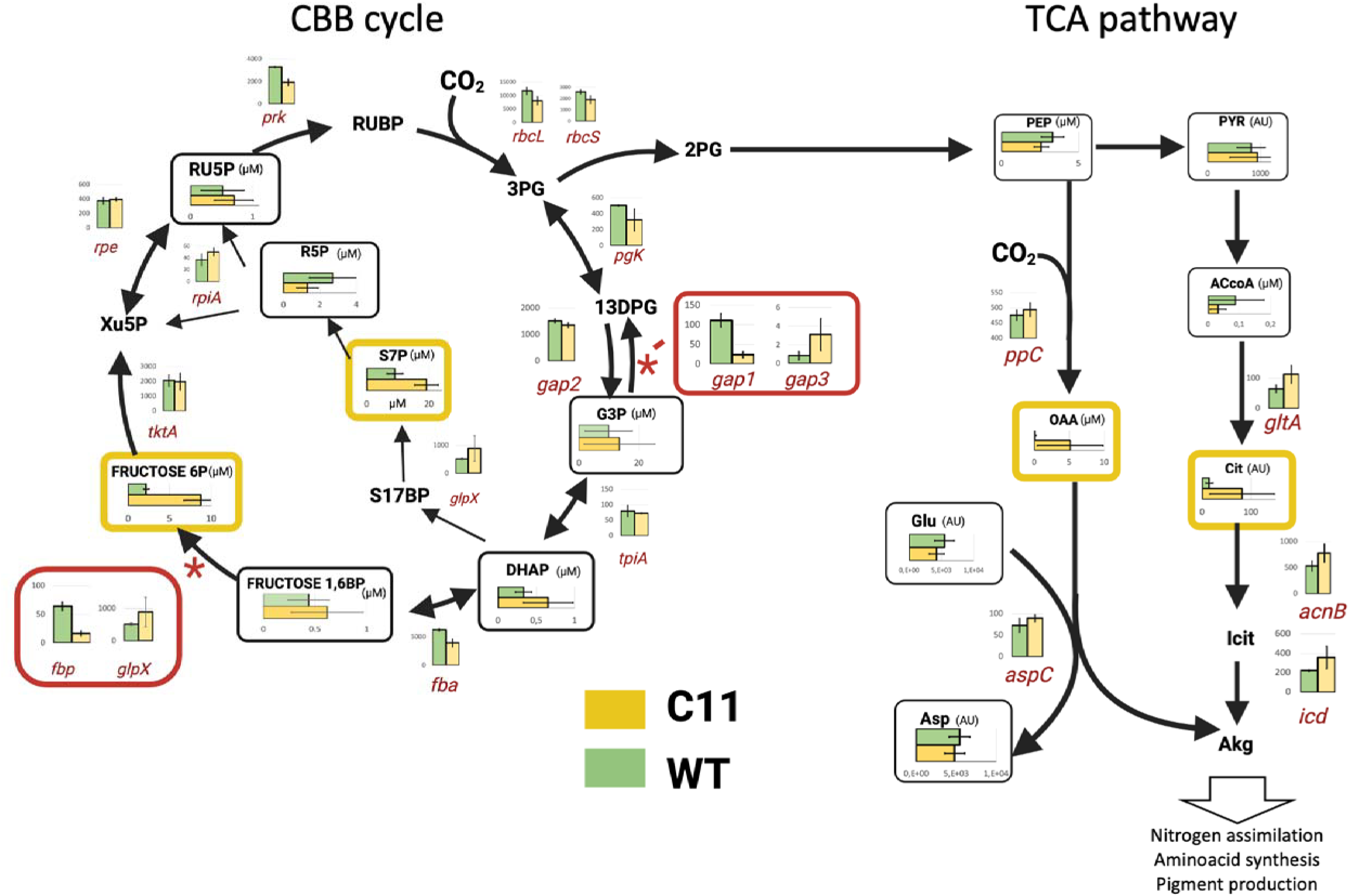
Diagram showing the levels of key metabolites and enzymes of the Calvin-Benson-Bassham (CBB) and Tricarboxylic Acid (TCA) cycle in the wt (green) and C11 (yellow). Vertical charts show the transcription levels of key enzymes in the pathway, as determined by transcriptomic analysis, indicated as FPKMs. Error bars indicate the standard deviation of three biological replicates. Horizontal charts show the levels of intermediate metabolites, as determined by metabolomic analysis as concentration (µM) or arbitrary units (AU). Error bars indicate the standard deviation of three independent experiments. Values for each enzyme/metabolite can be found in Supplementary table 2 for enzymes and Supplementary Table 3 for metabolites.

### C11 PRESENTS PERTURBATIONS IN THE CIRCADIAN CYCLE

The *sasA*Δ30 mutation caused a deletion in the kinase domain of SasA, a protein that phosphorylates the circadian regulator RpaA to its active conformation. Since an active RpaA is essential to sustain circadian transcription, this mutation could disrupt the circadian rhythm of C11. To test this end, we measured the expression of *sigC,* a prototypical Class I gene^32^. For this purpose, a yellow fluorescent protein (YFP_LVA) was transcriptionally fused to the P*_sigC_* promoter^32^. This transcriptional reporter was introduced into the NS1 neutral site of the wt, C11, *sasA*Δ30, P*aroK* and *hMT* strains. The expression profile was determined by measuring YFP expression in single cells using time-lapse microscopy, as described in Materials and Methods. Results are shown in Figure 4 and Supplementary Videos 1-4. As shown in Figure 4A, the fluorescence profile obtained from the wt showed the behavior expected for a prototypic Class I promoter, with a peak at dusk and a trough at dawn. This cyclic behavior was absent in C11, which showed arrhythmic YFP expression (Figure 4B). Among the point mutants, both *hMT* (Supplementary Figure 4) and P*aroK* (Figure 4D) showed robust circadian profiles, and the latter exhibited a 2-fold increase in amplitude compared to the wt. The arrhythmic phenotype was also observed in *sasA*Δ30 (Figure 4C), indicating that this mutation is key to the loss of circadian expression in C11. The defect in C11 rhythmicity was confirmed using a P*_psbAI_*-YFP construction, which yielded similar results (Supplementary Figure 5). Overall, results showed that C11 displayed an abnormal circadian cycle, with *sasA*Δ30 as the driving mutation.

**Figure 4.-.**
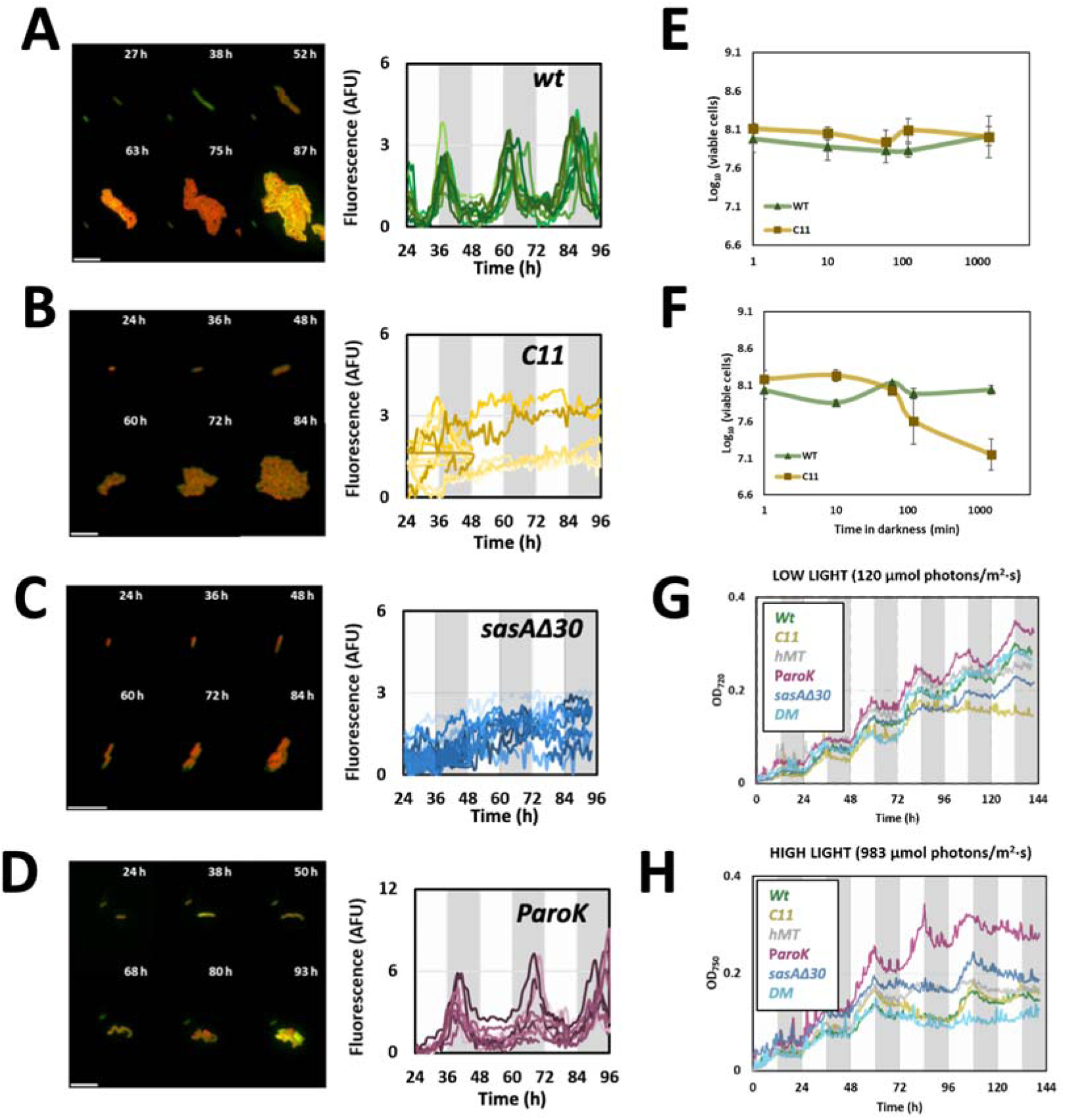
Circadian rhythm in C11 and individual mutants. (A-D). Microphotographs (left) and fluorescence traces (right) obtained from a P*_sigC_*_YFP transcriptional fusion in each of the strains indicated. Time-lapse experiments were performed on cultures pre-synchronized after growing for 72 h in LD conditions. Each trace on the graphs represents the values obtained for a single cell, tracked for 72 h. Dark and light areas of the charts indicate, respectively, the periods of subjective night and day. (E-F) Darkness-induced lethality. Cultures of the wt (green) and evolved (yellow) strains were grown under continuous low (E, 30 µmol photons m^-2^ s^-1^) or medium (F, 320 µmol photons m^-2^ s^-1^) light intensities and exposed to total darkness for a period indicated in the x axis. The number of viable cells after darkness exposure (y-axis) was determined by plating, as indicated in materials and methods. Each data point represents the average and standard deviation of three independent experiments. (G-H) Growth curves, monitored as OD720 (y-axis) along time (x-axis) under light / dark conditions. Cultures were grown under 0.04% CO_2_ (LC) concentrations and a regime of 12 h light + 12 h darkness (grey areas). Light intensity was set at 120 µmol photons m^-2^ s^-1^ (G) or 983 µmol photons m^-2^ s^-1^ (H). Under any of these conditions, C11 (dark yellow line) was unable to grow.

Circadian mutants like Δ*rpaA* strains, suffer from darkness-induced lethality, dying after short periods of time in the absence of light^15^. Since C11 exhibited circadian defects, we tested its sensitivity to darkness. For this purpose, we grew cells under continuous light, exposed them to a period of total darkness, and then measured the viability of the culture. As shown in Figure 4E, we observed no darkness-induced lethality when cells were grown at low light intensity (30 µmol photons m^-2^ s^-1^). When cells were grown at 320 µmol photons m^-2^ s^-1^, however, C11 showed a sharp decrease in viability after 60 minutes of darkness. Another characteristic phenotype of Δ*rpaA* circadian mutants is their inability to grow in light/dark cycles. We tested for this possibility in C11, and obtained growth curves in diel cycles of 12 h light/12 h darkness. Results, shown in Figure 4G-4H indicated that C11 was able to grow in these conditions, yet significantly slower than the wt.

### GENE EXPRESSION IN C11 IS CORRELATED TO A **Δ***rpaA* MUTANT AND UTEX 2973

C11 showed some hallmarks of a Δ*rpaA* strain, such as an arrhythmic expression in P*_sigC_* and P*_psbAI_*, darkness-induced lethality, and the repression of glycogen and pentose-phosphate pathways. For these reasons, we compared the transcriptomes of C11 (and each of the point mutants) with those of a Δ*rpaA* strain. For every gene and strain, we calculated the fold-change with respect to the wt. We then plotted these values against those of a Δ*rpaA* strain (Figures 5A-C). In the case of P*aroK* (Figure 5C), values scattered along a flat line across the x axis, indicating no correlation with Δ*rpaA* (Pearsońs r=0.06). However, a positive correlation was observed in C11 (r=0.89) and *sasA*Δ30 (r=0.71), as shown in Figures 5B and 5C. Genes highly repressed in these strains were also repressed in a Δ*rpaA* background. All 20 genes that showed the highest downregulation in C11 and *sasA*Δ30 were repressed by >5 fold in ΔrpaA (Figure 5E). Altogether, these results indicate that the *sasA*Δ30 mutation caused transcriptomic changes similar to that observed in a ΔrpaA mutant, suggesting a perturbed SasA-RpaA transduction pathway in *sasA*Δ30 and C11.

**Figure 5.-.**
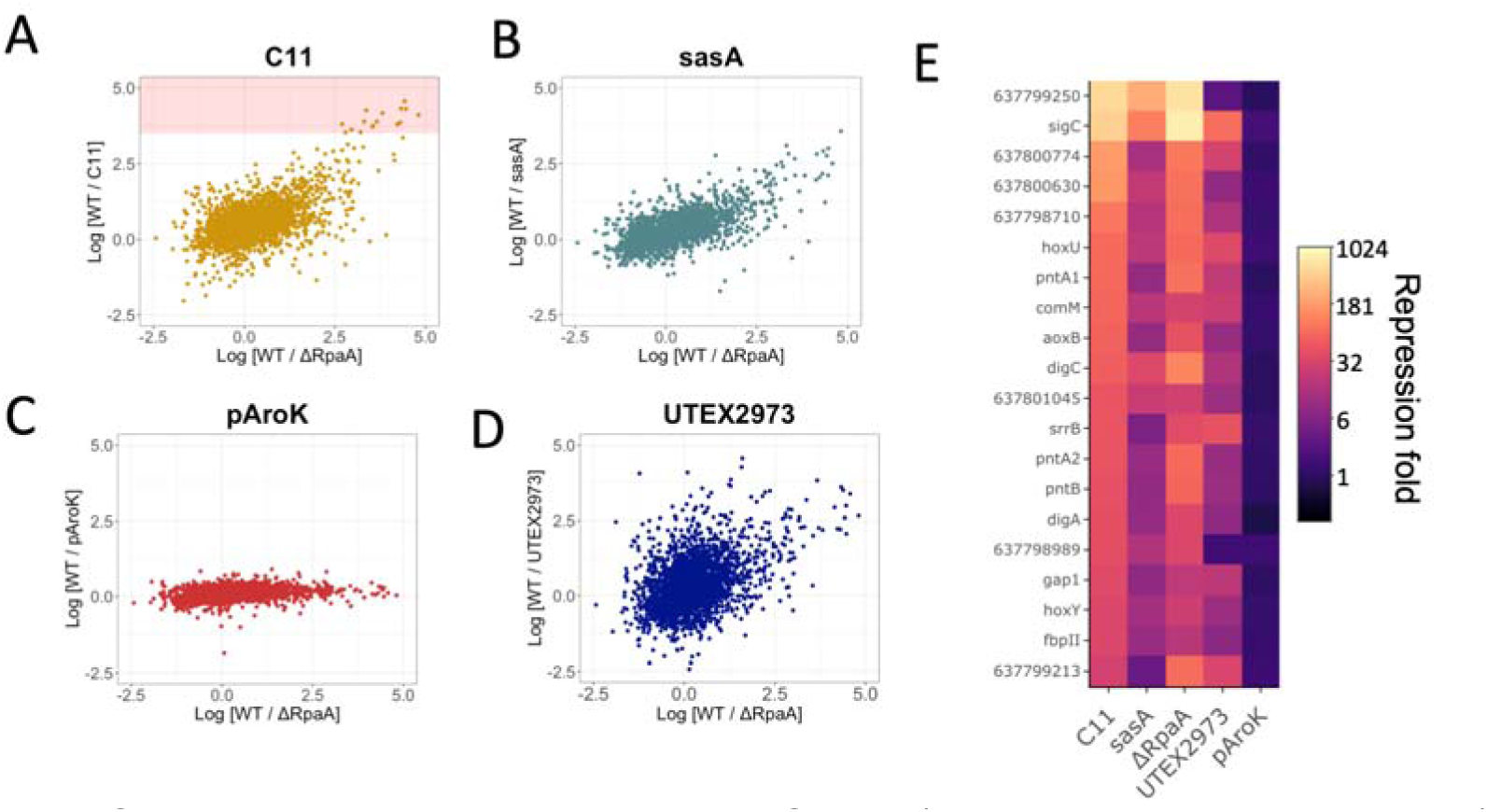
Correlation between the transcriptomes of C11, *sasA*Δ30, P*aroK*, UTEX 2973 and Δ*rpaA*. (A-D) Scatterplots showing the repression fold, expressed as the Log_10_ of the FPKMs exhibited by a particular gene in the wt, divided by the corresponding mutant (y-axis), compared to those of ΔrpaA (x- axis). The red area in panel 5A represents the levels corresponding to the 20 most repressed genes in C11. (E) Heatmap showing the repression fold, calculated as before, and expressed in the color chart shown in the legend, for the 20 most repressed genes in C11, in each of the mutants shown in the x-axis of the figure.

Intriguingly, UTEX 2973, a strain that, like C11, grows significantly faster under HL conditions, also harbors mutations in RpaA that are essential for its phenotype^23^. We thus took advantage of the transcriptomic data available for UTEX 2973, and compared it, as in the previous case, with that of a Δ*rpaA* mutant. As shown in Figure 5D, we found a positive correlation, albeit less marked than that of C11 and *sasA*Δ30 (r=0.25). However, when we analyzed the 20 genes with the highest repression in C11, 18 of them showed at least a 5-fold repression in UTEX 2973 (Figure 5E). These results suggest that mutations in the circadian cycle may be key to achieve a fast-growing phenotype.

### CHANGES IN THE PHASE AND AMPLITUDE OF CLASS I AND CLASS II GENES

Since C11 exhibited traits similar to a Δ*rpaA* mutant, we studied the circadian expression of its entire genome. For this purpose, we grew C11 and the wt for four days, in cycles of 12 h light/ 12h darkness. Total RNA was extracted at the dusk of the third day, and the dawn and dusk of the fourth. We then performed a transcriptomic analyses, comparing gene expression levels at these three time points. In the wt, a majority of genes (2159) exhibited a peak during dusk, thus behaving as Class I (Figure 6). A second set (428) showed a peak during dawn, characteristic of Class II genes. Finally, a total of 128 genes peaked neither at dusk nor dawn, and were considered non-circadian. We restricted our analysis to genes with a strong circadian character, selecting those in which the amplitude of the fluctuation was higher than four standard deviations of the average (p > 0.001). Using this criterion, we identified a total of 479 representative genes belonging to Class I (Figure 6A), and 43 genes belonging to Class II (Figure 6B). When we analyzed these genes in C11, we found profound changes in their expression profiles. The most apparent was observed in Class II genes, which dramatically changed their phase. Most of them showed a peak at dusk, indicating that, in C11, they behaved as typical Class I genes. This radical inversion of phase was observed in the entire class, affecting canonical genes like *purF*^9^. This effect was caused by a transcriptional repression during the night, which pushed transcription levels at dawn below the ones observed at dusk (Figure 6E).

**Figure 6.-.**
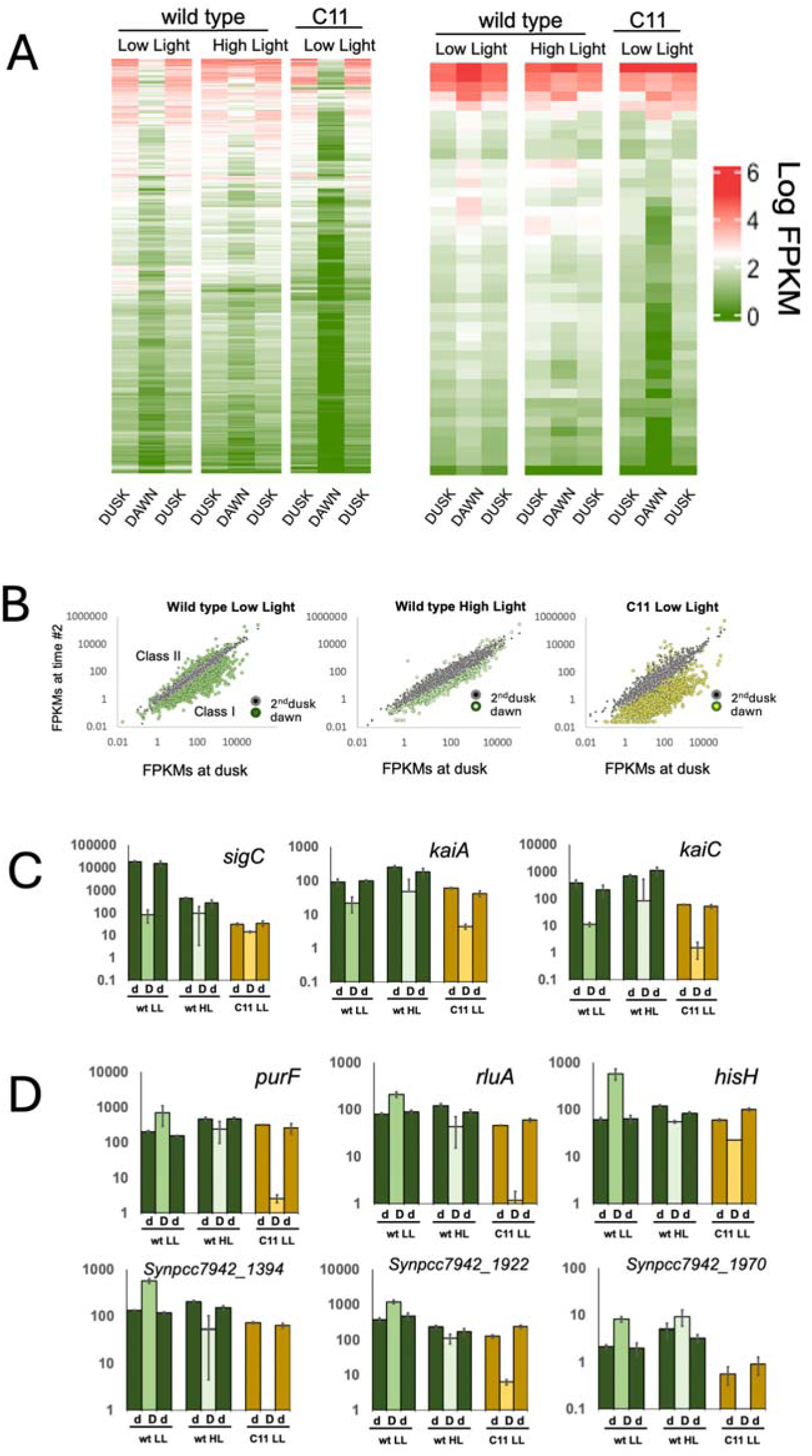
Perturbations in the circadian cycle. Heatmap showing the expression levels of Class I **(A)** and Class II **(B)** promoters, indicated as Log_10_ FPKMs. All cultures were grown in a regime of 12 h day/12 h night cycles for four consecutive days. The first column corresponds to the wt grown at a light intensity of 120 µmol photons m^-2^ s^-1^ (low light). The second column corresponds to the wt grown at a light intensity of 983 µmol photons m^-2^ s^-1^ (high light). The third column corresponds to C11 grown at a light intensity of 120 µmol photons m^-2^ s^-1^ (low light). For each column, the first lane corresponds to expression levels measured in the dusk of day 3, and the second and third lanes correspond, respectively, to the dawn and dusk of day 4. **(C)** Correlation of the expression levels of each gene at dawn and dusk. Cells were measured on the dusk of the first day (x-axis), dawn (coloured dots) and the dusk of the second day (black dots).Expression levels measured at the two sequential dusk periods are expected to be similar, thus falling in the diagonal of the image. Class I genes show higher expression during dusk, and should appear below the diagonal, while Class II genes appear above. The charts correspond, from left to right, to values obtained in the wt grown at 120 µmol photons m^-2^ s^-1^ (LL), the wt grown at 983 µmol photons m^-2^ s^-1^ (HL), and C11 grown at 120 µmol photons m^-2^ s^-1^ (LL). **(D)** Transcription levels for selected Class I genes, including *sigC* (left), *kaiA* (middle) and *kaiC* (rightmost panel). Bars correspond to transcription levels observed at dusk of day 3 (d), dawn of day 4 (D) and dusk of day 4 (d). Error bars indicate the standard deviation. **(E)** Transcription levels for selected Class II genes. Bars and legend as in (D).

In C11, Class I genes also experienced decreased transcription during nighttime. However, since these genes are naturally downregulated during the night, the phase of this class was generally unaffected, and the global effect was an increase in the amplitude of the circadian fluctuation. We nevertheless identified a subset of Class I genes in which the downregulation of transcription also occurred during daytime, leading to the loss of their circadian character. This was the case of *sigC* (Figure 6D), confirming the results obtained in time-lapse experiments (Figure 4B). Other classical Class I genes, such as *kaiA* or *kaiC* were, however, unaffected and maintained the phase and amplitude of their circadian fluctuation (Figure 6D). Results thus demonstrated that the circadian cycle is severely altered in C11, but not entirely abolished.

The circadian alteration in C11, together with its constitutive repression of genes downregulated by high light, suggested that light intensity during the day could somehow influence circadian rhythms. To study this possibility, we analyzed the impact of high intensity illumination during daytime in the circadian cycle of PCC 7942. We repeated the previous experiment, growing the wt in diel cycles with the light intensity during daytime increased to 983 µmol photons m^-2^ s^-1^. Results, shown in Figure 6A and B, indicated a global nighttime repression of all Class II genes, which again led to the reversal of their circadian phase. For Class I genes, however, expression levels during the night were elevated, leading to shallower fluctuations. In *sigC,* and the subset of genes that were downregulated and arrhythmic in C11, the circadian cycle was also abated. Thus, C11 presented some circadian perturbations that were also observed in wt cells exposed to high intensity illumination. The switching of Class II genes to a Class I phase, and the downregulation and loss of rhythmicity in *sigC* were the most conspicuous features.

## DISCUSSION

Cyanobacteria are key primary producers in the biosphere, yet the environmental and genetic factors constraining their growth are poorly understood^33^. Strains from the same species exhibit radically different reproductive rates and environmental preferences^23^. To study the causes of this phenotypic diversity, we performed an LTE experiment, growing the wt strain for 1200 generations under intense, continuous illumination. The resulting population, C11, showed a 600% increase in growth rate. In the classical LTE experiment of Richard Lenski, it took *E.coli* 50,000 generations to achieve a 70% growth rate, indicating that, in *S.elongatus,* adaptation was faster and the fitness gain much higher^34^. The recent isolation of UTEX 2973, a strain with a fast-growing phenotype similar to C11, prompted the question of whether fast growth could be actually a wt feature of *S.elongatus*, lost in PCC 7942 due to laboratory domestication^35^. Two lines of evidence, however, suggest otherwise. On the one hand, UTEX 2973 and C11 show no mutations in common, and genes essential for fast growth in C11 (such as *aroK*) are not mutated in UTEX 2973. On the other hand, recent isolates of *S. elongatus* grow at rates and environmental conditions similar to PCC 7942 and other legacy strains, suggesting that the fast-growing phenotype is not a lost common trait, but a specific adaptation^22^. Like other freshwater β-cyanobacteria, *S. elongatus* is primarily found in ponds and small water reservoirs prone to large and sudden environmental perturbations^36^. In such contexts, a high phenotypic plasticity may contribute to cyanobacterial survival. The fact that the fitness gain experienced by C11 is entirely contingent on the environmental conditions of the LTE (Figure 1A), suggests that phenotypic plasticity may be a way for the cyanobacterium to fine-tune its internal state to the particular ecological constraints of the environment.

Although C11 and UTEX 2973 do not show mutations in common, both strains converged to similar transcriptomic states, suggesting perturbations in the SasA-RpaA axis. In C11, these defects were caused by the c-terminal deletion of SasA, resulting in a profound alteration of the circadian cycle. Suppressing the expression of Class II genes and the night metabolism may be beneficial in continuous light, as it has been previously hypothesized for cyanobacterial strains with no clock^37^. Since the cycle is endogenously generated by the PTO, cells experience circadian transcription even when under perpetual daylight. Thus, abolishing the subjective night phase, a predominantly catabolic period when cell division is gated, may increase the overall growth rate. Data from C11 and UTEX 2973, however, indicate that while mutations in SasA-RpaA are necessary, they are not sufficient for a fast-growth. This agrees with previous results from Δ*rpaA* mutants, which showed no observable fitness advantage under continuous illumination^38,39,40^. Both C11 and UTEX 2973 required secondary mutations, probably to funnel the metabolic potential liberated by the abatement of night metabolism into effective biomass production. In C11, increased levels of metabolic intermediates of the CBB cycle were observed, and the changes in the expression levels of *gap1*/*gap2* isoenzymes indicated a downregulation of catabolic pathways. Whether this was caused by alterations in the circadian cycle or by secondary mutations remains to be elucidated. The role of the P*aroK* mutation in the phenotype of C11 may be related to stress tolerance. In *Synechococcus* and *Synechocystis* mutations in AroK improve growth in high light and high temperature, but the exact molecular mechanism through which this mutation contributes to fitness remains uncertain^41^. There are no mutations in the shikimate pathway in UTEX 2973, thus it is unlikely that both strains achieved fast-growth through identical mechanisms. This opens the question of whether combining the mutations of both strains may lead to an additive effect. Such strategy could lead to the generation of faster-growing strains, useful chassis for biotechnological and synthetic biology applications.

In *Synechococcus*, fluctuations in the PTO are converted into a transcriptional oscillation through RpaA, but complex downstream regulation decomposes this initial tempo into transcriptional cycles of different phase and amplitude^42,43^. Our results demonstrate that light intensity specifically modulates these cycles, and that adaptation to high intensity illumination requires mutations in the circadian pacemaker. Previous work using computational models predicted that the fitness benefit conferred by the clock would decrease with growing light intensities^37^. Our results support this prediction, suggesting, in turn, that the circadian rhythms of UTEX 2973 are probably perturbed too. Altogether, the phenotypic and transcriptomic characterization of C11 indicate that, in cyanobacteria, the circadian rhythm is not a mere instrument to separate day and night physiology. It is also an adaptive mechanism, providing phenotypic plasticity to ensure growth in different light intensities and environmental conditions.

## MATERIALS AND METHODS

### Cyanobacterial strains and culture growth conditions

*Synechococcus elongat*us PCC 7942 (PCC 7942) and all derived strains were routinely grown and maintained in liquid BG11 medium supplemented with appropriate antibiotics. Antibiotics used for selecting PCC 7942 were neomycin at 5 or 25 µgml^-1^ (Neo5 or Neo25), spectinomycin at 10 or 20 µg ml^-1^ (Sp10 or Sp20) and streptomycin at 10 or 50 µg ml^-1^ (Sm10 or Sm50). Except otherwise stated, cells were grown under a continuous light flux of 60 µmol photons m^-2^ s^-1^ from white fluorescence lamps, in a Sanyo Plant Growth Chamber, at 30°C and constant ambient air supply. Experiments with different light intensities, diel cycles and CO_2_ concentrations were performed in a MC 1000-OD Multicultivator (Photon Systems Instruments), equipped with programmable temperature, CO_2_ and light intensity (cold white LED). Doubling times were determined from growth curves performed in a MC 1000-OD Multicultivator (Photon Systems Instruments). Growth was estimated from OD_720_ and OD_680_ measurements obtained every 10 minutes by the multicultivator. The effective growth rate (K’) was obtained by the slope of the LogOD_720_ in the early exponential phase of each growth curve. The doubling time was calculated as Ln2/K’

### Experimental evolution

For the evolution experiment, a population of PCC 7942 was grown in liquid BG11 medium in 200 ml glass tubes with a continuous light flux of 1,313 µmol photons m^-2^ s^-1^ at 41°C and 5% of CO_2_. A control population was grown in parallel, at 25°C, atmospheric CO_2_ levels and a continuous light flux of 65 µmol photons m^-2^ s^-1^. Cultures were refreshed every 24 generations through a 1:130 dilution. During the long-term evolution experiment, the progressive increase in fitness was monitored by the time required for the culture to achieve the 24 generations. Whole genome sequencing was performed at generations 0, 824 and 1,284 to track the emergence of mutations during the experiment.

### Genetic constructions

Genetic engineering in PCC 7942 was performed through natural transformation. For each transformation, approximately 4×10^9^ cells, equivalent to 10 μg of chlorophyll, were mixed with 500 ng of DNA, and incubated in the dark for 16 h at 30°C and 100 rpm. Transformation mixtures were then deposited onto a 0.45 μm nitrocellulose filter (Millipore) and incubated for 24 h on top of BG11 plates supplemented with appropriate antibiotics, at 30°C and continuous light. The filter was transferred to new plates with antibiotics every 24-72 h. Transformation colonies appeared after 7-14 days. Segregation of mutants was achieved by repeatedly striking individual transformant colonies on selective plates. Mutant genotypes were confirmed by PCR and DNA sequencing using specific primers.

The gene replacement method used to insert the mutations from C11 in the wt strain was based on Matsuoka et al.^44^. Briefly, a streptomycin resistant wt strain (MSM1) was obtained by naturally transforming PCC 7942 with the K43R recessive allele of the ribosomal protein S12 (*rps12-R43*) and selection of Sm^R^ resistant transformants. This Sm^R^ strain was used for the introduction of the individual mutations of C11 in the wt background. A set of two plasmids was designed to insert each mutation. The first one contained a kanamycin/neomycin resistance cassette and the *Synechocystis* sp. PCC 6803 wt version of the *rps12* gene cloned under the promoter of PCC 7942 *psbA*I gene.

This gene was flanked by the regions immediately downstream or upstream the gene of interest. Transformants were selected on Neo25 plates and confirmed by their Neo^r^/Sm^s^ phenotype and PCR. To introduce the desired mutation, a second plasmid containing the mutated gene of interest and its flanking regions was introduced through natural transformation. These transformants were then selected through their Neo^s^/Sm^r^ phenotype. Transformants were checked through PCR and Sanger sequencing using appropriate primers.

### Darkness-induced lethality assays

The dark-induced lethality phenotypes were determined by measuring the number of viable cells surviving a pulse of total darkness. For this purpose, cells were grown under continuous illumination until reaching OD_720_ 0.4-0.6. At this moment, the light was shut down for a variable time (0, 10, 60, 120 or 1440 minutes). The survival rate of the culture after darkness was calculated by counting the number of viable cells on BG11 media obtained immediately before and after the pulse of darkness.

### Time-lapse microscopy

YFP-LVA cassettes transcriptionally fused to P*_sigC_* or P*_psbA_I* promoters were introduced in the neutral site 1 (NS1) of each of our strains through homologous recombination. These cassettes, taken from Martins et al, contained a Sp resistance gene for selection. To introduce the construction in C11, which has lost its natural transformation ability due to the *pilA* mutation, a mobilizable plasmid was constructed to shuttle the cassette by conjugation. For this purpose, a RP4 oriT was introduced in plasmid pLA35 from Martins et al, which contained, the P*_sigC_*_YFP-LVA and construction. This allowed the mobilization of the cassette to C11 from a MDFpir strain of *E. coli*, using the protocol described by Encinas et al. (2014)^45^. In the rest of the strains, the cassette was introduced through natural transformation following the procedure described above.

Time-lapse images were taken using a Nikon Eclipse Ti2 microscope. For this purpose, cells were grown in liquid BG11 for 2-3 days in 12 h light-12 h darkness conditions to synchronize the population. A 10 µL sample was loaded on a solid BG11-agarose (1.5%) pad and entrained in the microscope for another day. Illumination was achieved using the transmitted LED source under Khöler illumination at 387 – 775 PAR luxes. Light intensity was adjusted using a PM100A light power meter equipped with a slide- type probe. The correlation between PAR lux intensity and µmol photons m^-2^ s was calibrated using a LI-250A PAR-lightmeter (LI-COR^®^). After entrainment, the cells were kept in continuous illumination for 72 h. Brightfield, red fluorescence and yellow fluorescence images were taken every 45 minutes under a 60x/1.4 NA PLAN apochromat objective and an ORCA Flash4.0 (Hamamatsu^®^) camera with a 2×2 binning to reduce background noise. Green and red fluorescence was captured using 472/30nm excitation – 520/35nm emission and 578/21nm excitation – 641/75nm emission filter cubes (Semrock^®^) respectively. Timelapses were simultaneously recorded at five different positions separated at least two fields of view and disposed as the five faces of a dice array. Between different timepoints, transmitted illumination was maintained at the central position with the condenseŕs field diaphragm fully open, thus assuring equal light intensity during transmitted illumination for the different positions without cross- illuminating. During the duration of the experiment, cells were maintained in focus using Nikońs Perfect Focus System, which was let to stabilize for at least 10s at each position before image acquisition. To maintain transmitted illumination ON during timepoints and stage movements, we took advantage of the microscope softwarés (NIS elements) scripting capacity. The resulting fluorescent images were manually segmented with the software Oufti to determine the amount of green and red fluorescence present in the tracked cells along the time-lapse^46^.

### RNA-seq analysis

Total RNA extraction protocol was an adaptation from Hein et al.^47^ 30 ml of a culture at OD_720_ ∼ 0.6 were collected by centrifugation at 4000 *g*, 10 min RT. The pellet was resuspended in 1.5 ml of Trizol (Ambion-USA) and the suspension was divided in two pre-cooled RNAse-free eppendorf tubes and immediately frozen in liquid nitrogen and stored at −80°C. To extract the RNA, the samples were incubated at 65°C for 15 min, being vortexed several times in the process. 525 µl of chloroform:isoamiloalcohol (24:1) were added. After a 10 minutes incubation at RT, mixing gently several times, the samples were centrifuged at 6000 *g*, three minutes at RT. The aqueous phase (470 lµ) was transferred to new RNAse-free tubes and 470 µl of chloroform:isoamiloalcohol (24:1) were added, repeating the previous steps. The aqueous phase (370 µL) was transferred again to a new tube, mixed with the other half it was previously separated from- One volume (740 µl) of isopropanol was added, gently mixed and left precipitating overnight at −80°C. The RNA was pulled down by 12000 *g* centrifugation at 4°C, for 30 minutes. The pellet was washed three times with 200 µL of ethanol 75% and left to dry. The clean RNA was resuspended in 54 µl of RNAse-free distilled water. Two µl per lane were loaded in a 2% agarose gel to value sample’s integrity, and two µl were used to measure purity and concentration in a Nanodrop (Thermo). The remaining 50 µl were subjected to a DNAse digestion protocol (Invitrogen) and an additional cleaning step running the final volume through an RNA extraction kit column (Qiagen).

The sequencing reads from each sample were trimmed for Illumina adapters remnants and low-quality regions (<Q25) using Trim-Galore (v0.6.2)^48^. Surviving high-quality reads were posteriorly aligned against three bacterial rRNA databases (5S, 16S and 23S) from SILVA (https://www.arb-silva.de/) and Rfam (https://rfam.org/) with SortMeRNA to filter out rRNA fragments (v4.3.6)^49^. The processed reads were aligned against the complete genome sequence of *Synechococcus elongatus*, strain PCC 7942 (GenBank accession number: GCF_000012525.1) using Hisat (v2.2.1)^50^, processing the intermediate alignment files and merging the technical replicates with Samtools (v1.6)^51^. Individual sample gene expression levels were quantified with featureCounts (v2.0.1)^52^. Differential expression of mRNA transcripts was computed with DESeq2 (v1.34.0)^53^, using a padj value < 0.05 as statistical cutoff. In parallel, normalized expression values were calculated using StringTie (v2.2.1)^54^ in transcripts per million (TPM).

### Metabolomics preparation and analysis

The protocol followed for metabolome extraction was adapted from the one described by Prasannan et al. (2018)^29^. Briefly, a volume of 20 mL with an OD_720_ of 0.6 was withdrawn from the culture grown in the desired conditions and vacuum filtered through a Durapore® 0.45 µm PVDF membrane (Merck Millipore-Ireland). The filter with the biomass was immediately inverted into a petri dish with 1.6 ml of pre-cooled methanol for quenching and incubated at −80°C for an hour with periodic rubbing of the filter against the container to aid cell lysis. Subsequently, the solution was transferred to a reaction tube, and the filter was extracted with 2.4 ml of chloroform. Both fractions were merged and vortexed for 20 minutes (30 seconds on, one minute off in ice) to disrupt cells. After addition of two ml of H_2_O, the mixture was vortexed for 10 minutes.

After centrifugation (3,200 *g*, 15 minutes, 4°C), two aliquots (750 µl each) of the transparent upper phase were collected and vacuum dried for eight h. Samples were stored at −20°C. Upon analysis, samples were resuspended in 180 µl of H_2_O and mixed with 20 µl of internal standard and insoluble particles were removed by centrifugation (17,000 *g*, 10 min, RT). The internal standard consisted of extract from *E. coli* grown on U-^13^C-glucose. Metabolites were quantified using LC-MS/MS according to the method of McCloskey et al. (2016)^55^, and the chromatograms were analyzed using the MultiQuant™ software (Sciex, CA, USA). Metabolites were quantified against calibration curves of authentic commercial standards using the internal standard.

## List of Supplementary Materials

### Supplementary Figures

S. Figure 1- C11 grows slower than the wt at low CO_2_ concentrations.

S. Figure 2.- C11 has lost its natural transformation ability.

S. Figure 3.- Genes overexpressed in high light.

S. Figure 4.- Circadian rhythm in the *hMT* strain with a P*_sigC_*_YFP construction.

S. Figure 5.- Circadian rhythm in P*_psbAI_*_YFP constructions.

S. Figure 6.- Viability after dark pulse in the single mutants.

### Supplementary Videos

S. Video 1- P*_sigC_*-YFP time lapse in the wt.

S. Video 2- P*_sigC_*-YFP time lapse in C11.

S. Video 3- P*_sigC_*-YFP time lapse in P*aroK*.

S. Video 4- P*_sigC_*-YFP time lapse in *sasA*Δ30A.

### Supplementary Tables

S. Table 1- Genes mutated in the LTE.

S. Table 2- Transcriptomic data for growth in LL.

S. Table 3- Metabolomic data.

S. Table 4.- Transcriptomic data for growth in LD.

S. Table 5- Strains used in this work.

S. Table 6- Plasmids used in this work.

## ACKNOWLEDGEMENTS

The authors wish to thank Prof. James Locke from Sainsbury Laboratory for kindly providing us with the plasmids that allowed the measurement of pSigC expression in single cells. Genomic and transcriptomic analyses shown here were performed with the help of the National Center for Genomic Analysis (CNAG). This work was funded by grants PID2019-110216GB-I00 and TED2021-130689B-C31 from the Spanish MICN/AEI and European Next Generation funds to RFL and grants NNF10CC1016517 and NNF18CC0033664 from The Novo Nordisk Foundation to P.I.N.

---

**Supplementary Figure 1.-.**
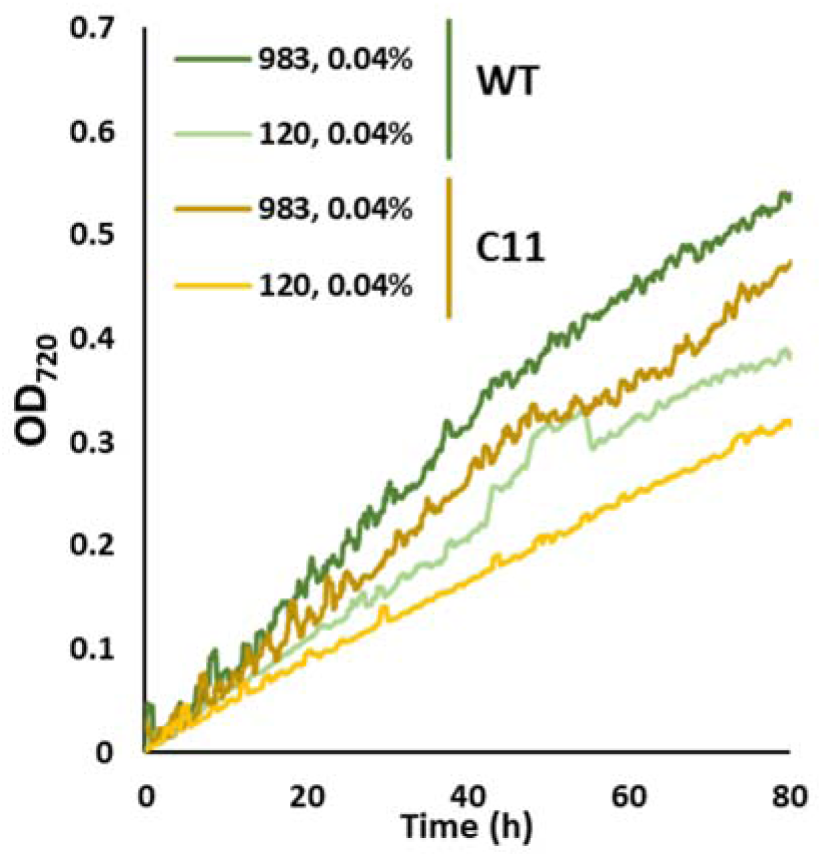
The evolved strain grows slower than the wt at low CO_2_ concentrations. Growth curves corresponding to the wt (green lines) and the evolved strain (yellow lines), measured as OD_720_ (y-axis) across time (x-axis). All measurements were performed in BG11 medium at HT, LC (41°C, 0.04% CO_2_) and the light intensities shown in the chart (HL: 983 µmol of photons m^-2^ s^-1^, LL:120 µmol of photons m^-2^ s^-1^).

**Supplementary Figure 2.-.**
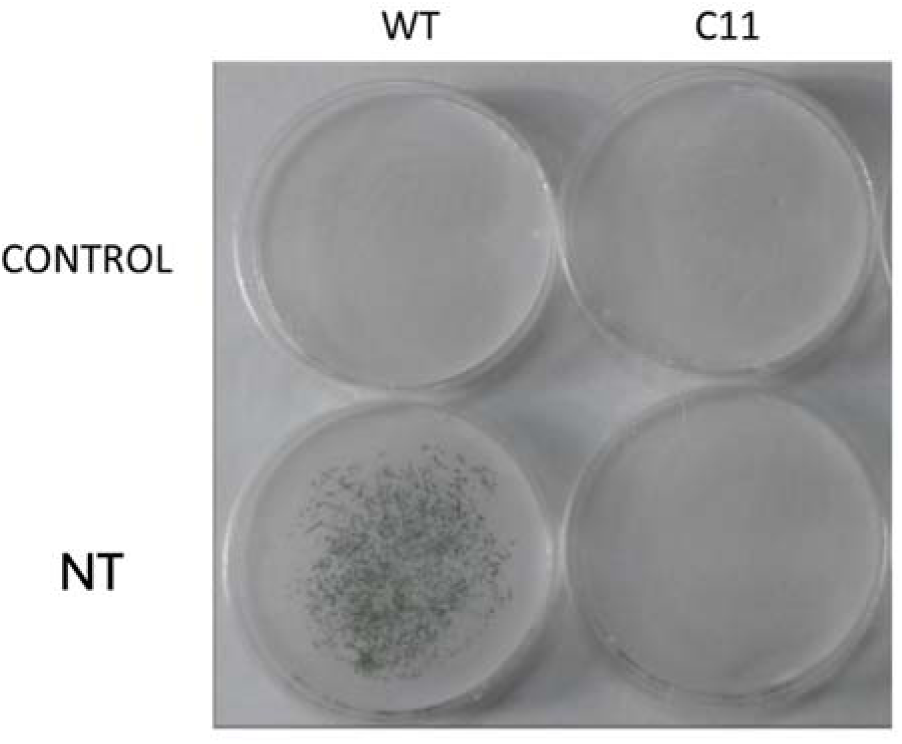
Natural transformation of the wt and the evolved strains. Control with no DNA and natural transformation (NT) with the plasmid pMSM2 of the wt (left) and the evolved strain, C11 (right).

**Supplementary Figure 3.-.**
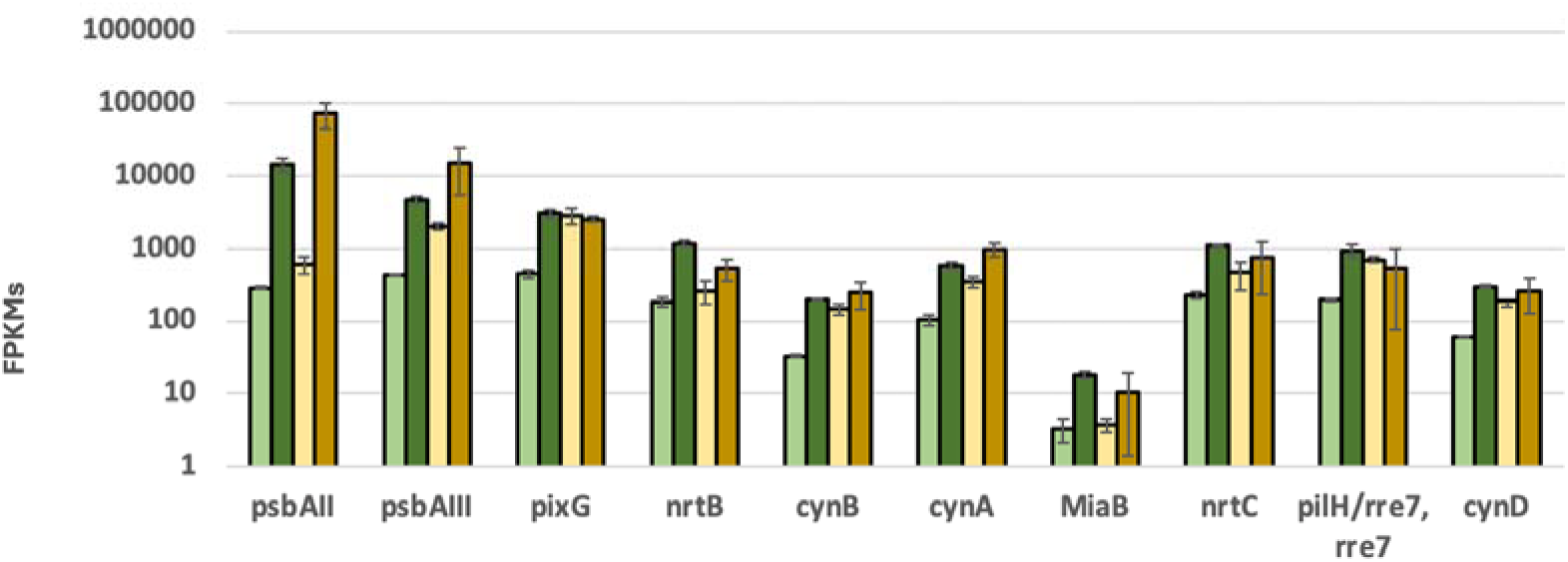
Gene expression levels of genes activated during growth in high light. The bars represent the average and standard deviation of three separate experiments.

**Supplementary Figure 4.-.**
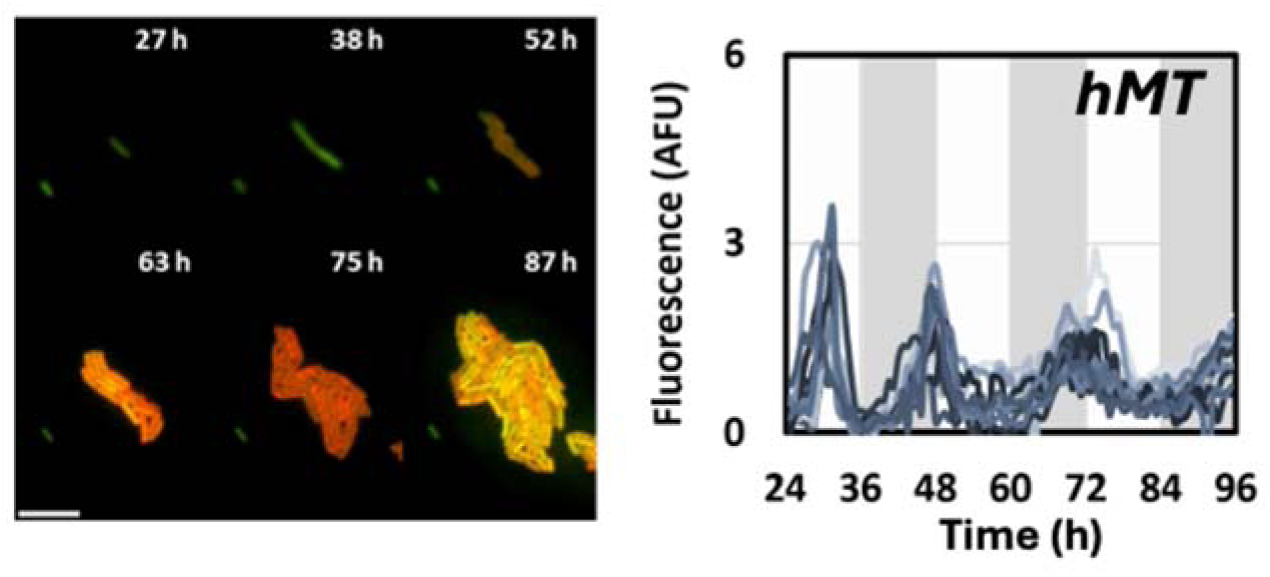
Circadian rhythm of *sigC* in the *hMT* mutants. Microphotographs (left) and fluorescence traces (right) obtained from a P*_sigC_*_YFP transcriptional fusion in the *hMT* strain.

**Supplementary Figure 5.-.**
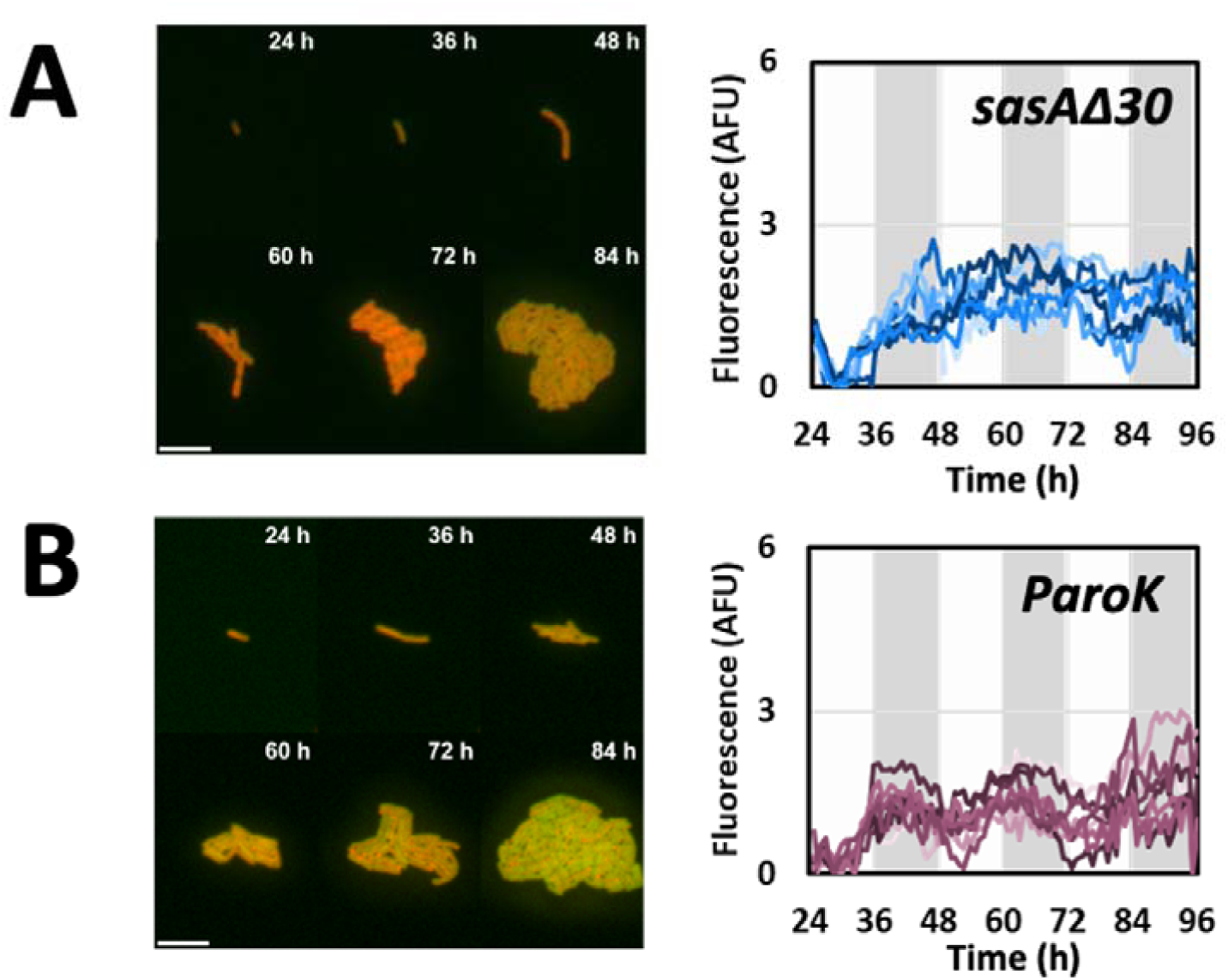
Circadian rhythm of psbA1 in individual mutants. Microphotographs (left) and fluorescence traces (right) obtained from a P*_psbA1_*_YFP transcriptional fusion in the *sasA*Δ30 mutant (A) and the P*aroK* mutant (B)

**Supplementary Figure 6.-.**
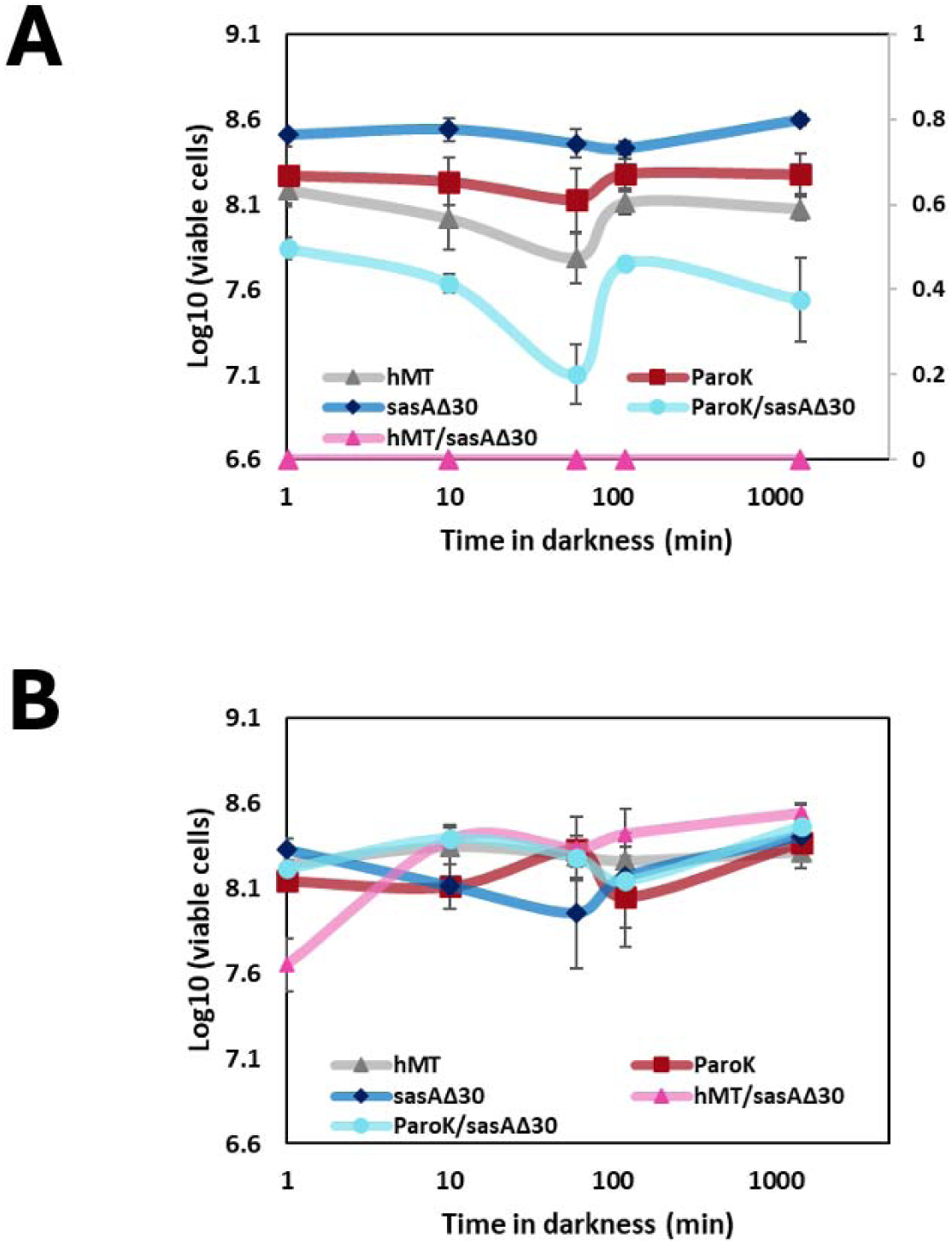
Viability after dark pulse in the single mutants. Cultures of the individual and double mutants were grown under continuous low (A, 20 µmol photons m^-2^ s^-1^) or high (B, 320 µmol photons m^-2^ s^-1^) light intensities and exposed to total darkness for a period indicated in the x axis. The number of viable cells after darkness exposure (y axis) was determined by plating, as indicated in materials and methods.

